# Electrical stimulation precisely reproduces naturalistic spiking activity in complete and intermixed neural populations in the primate retina

**DOI:** 10.64898/2026.05.04.721188

**Authors:** A.J. Phillips, Amrith Lotlikar, Alexandra Kling, Madeline Hays, Praful Vasireddy, Bella Hofflich, Aviv Sharon, Michael A. Sommeling, Jeff Brown, Claire Baum, Huy Nguyen, Ye-Lim Lim, Pawel Hottowy, Alexander Sher, Alan M. Litke, E.J. Chichilnisky

**Affiliations:** Department of Electrical Engineering, Stanford University, Stanford CA 94305, USA; Department of Neurosurgery, Stanford University, Stanford CA 94305, USA; Department of Ophthalmology, Stanford University, Stanford CA 94305, USA; Department of Bioengineering, Stanford University, Stanford CA 94305, USA; Hansen Experimental Physics Laboratory, Stanford University, Stanford CA 94305, USA; AGH University of Krakow, Faculty of Physics and Applied Computer Science, 30-059 Krakow, Poland; Santa Cruz Institute for Particle Physics, University of California, Santa Cruz, CA 94064, USA

## Abstract

Neurostimulation technologies have the potential to restore complex brain functions by evoking naturalistic patterns of neural activity. However, faithfully mimicking the brain’s natural neural code in diverse neuron types remains an unsolved challenge. Here we exploit focal electrical stimulation and recording to reproduce natural visual signals at cellular resolution in two intermixed and oppositely-tuned neural populations in the isolated macaque retina. We calibrated and delivered electrical stimulation sequences to precisely mimic recorded visually-evoked responses to natural images in complete populations of ~50 ON and OFF parasol cells uniformly sampling ~4° x 4° of visual angle. Distinct and opposing visually-evoked response patterns in the two intermixed populations were closely replicated spike-by-spike in 5 retinal preparations using electrical stimulation, reproducing spike trains with 51 ± 17% lower variability than repeated presentations of the same visual stimulus. Focal electrical stimulation also replicated traveling waves of neural activity in response to visual motion, and highly structured artificial response patterns. These findings demonstrate for the first time that neurostimulation can match or exceed biological signaling fidelity in large, complete and intermixed neural populations.

## Introduction

To explore and restore vital brain functions, neural interfaces of the future must be capable of reproducing the *neural code* — precise, coordinated spatiotemporal patterns of electrical activity in diverse types of neurons. Recent advances in all-optical approaches combining optogenetic actuators with genetically encoded activity sensors have made progress toward this goal, enabling novel experiments to interrogate and manipulate the function of neural circuits in animal models [1–9]. However, all-optical approaches require delivery and expression of transgenes in specific target cell populations, a significant technical hurdle that limits scientific and translational potential. Electrical recording and stimulation, on the other hand, are well-established for both preclinical science and clinical translation [10–15], but have yet to demonstrate precise control of large, diverse neural populations. Because of these obstacles, the promise of neurostimulation technologies for reproducing neural codes remains unfulfilled. Epiretinal implants for vision restoration are one important example. The neural code of the retina consists of distinctive patterns of activity in more than 20 retinal ganglion cell (RGC) types, each of which uniformly covers the visual field and projects to distinct targets in the brain (see [16]). In particular, multiple ON and OFF RGC types contribute in a push-pull manner to visual perception and must be stimulated precisely and independently to avoid sending conflicting information about light intensity to the brain. Unfortunately, existing epiretinal implants indiscriminately activate the intermixed RGCs of various types, failing to recreate the natural visual representation [17]. To reproduce the neural code of the retina, single-cell and single-spike control of multiple distinct neural populations simultaneously will likely be required.

Here we develop and test such an approach for the first time. We use large-scale multi-electrode stimulation and recording — a laboratory prototype of a future bidirectional epiretinal implant — to evoke naturalistic patterns of activity in two distinct and intermixed neural populations in the primate retina *ex vivo*. Dozens of ON and OFF parasol cells (two of the four numerically dominant RGC types in primates) completely covering a region of visual space were identified and targeted for recording and stimulation. Electrode and current level combinations yielding selective stimulation of each cell were identified and then exploited in the same experimental session to create optimized spatiotemporal stimulation sequences for both populations simultaneously. This approach accurately replicated recorded responses to natural images in the two cell types, decoupling their highly differentiated responses and thus creating a naturalistic visual signal. Traveling waves of activity in response to moving objects and highly structured responses not resembling the natural visual code were also accurately reproduced, further validating the approach as a tool for vision restoration and scientific investigation.

## Results

To reproduce the neural code of the primate retina in two distinct and intermixed neural populations, electrical stimulation and recording were performed using a high-density 512-electrode array system. Recorded responses to white noise visual stimuli were used to identify locally complete populations of two major retinal ganglion cell (RGC) types – ON and OFF parasol cells – and electrical stimulation and recording were used to calibrate the effect of current delivered through each electrode on each cell. Next, responses to natural images and moving bars were recorded in the two complete populations and then used to identify the target neural code to be replicated. Finally, in the same experimental session, the calibration was used to deliver current in spatiotemporal patterns optimized to replicate those target responses, and the fidelity of the replication was evaluated. The calibration was also used to evoke highly structured but artificial patterns of activity in the two cell types.

### Bidirectional electrical targeting of two interspersed neural populations

MEA recordings during white noise visual stimulation were spike-sorted (see Methods) and used to compute the spike-triggered average (STA) stimulus for each recorded cell in the preparation. Features of the STA and spike train autocorrelation function were then used to identify ON and OFF parasol cells [18–20]. The receptive fields of each cell type formed a uniform mosaic covering the recorded region, confirming cell type identification and indicating nearly complete sampling of the two populations in the region of retina recorded (Fig. 1A). The electrical image of each cell (average voltage waveform of its spikes on all electrodes) was also obtained (Fig. 1B, left), and was later used to identify electrically-evoked spikes (see Methods).

**Figure 1.**
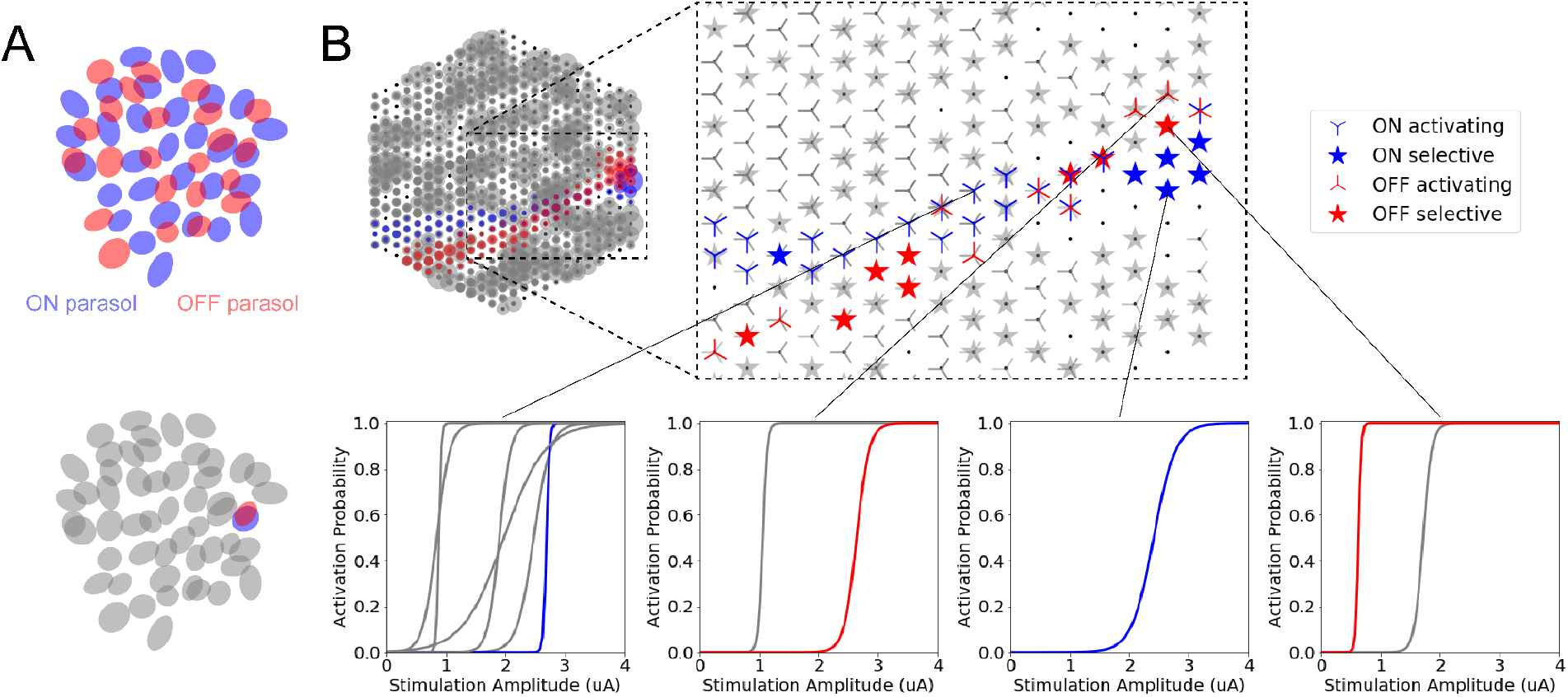
Bidirectional visual and electrical stimulus calibration. (A) ON and OFF parasol RGC receptive field mosaics. The ellipses denote the 2σ boundary of a 2D Gaussian fit to the spatial component of the STA. Bottom: A highlighted case of highly overlapping ON and OFF parasol RGCs. (B) Left: Superimposed spatial electrical images (EIs) of all ON and OFF parasol RGCs. The collection of blue (red) points denote the electrical image of the highlighted ON (OFF) parasol RGC, with the size of each point proportional to the peak recorded voltage on that electrode during the spike of that cell. The collection of gray points denote the electrical images of all other parasol RGCs. Right: Electrical response characteristics of the parasol RGCs in response to stimulation at individual electrodes. The collection of blue (red) markers denote the activating and selectively activating electrodes for the highlighted ON (OFF) parasol RGC. The activation curves at several stimulating electrodes are shown.

Single-electrode stimulation was then used to determine how current passed through each electrode influenced the firing of each recorded parasol cell: at each of the 512 electrodes, triphasic pulses of total duration 150 µs were delivered, with current amplitude 0.1-4 µA, and 15 trials at each current amplitude. The recorded activity immediately after electrical stimulation was fed through a custom real-time spike sorter to identify directly-evoked spikes in the ON and OFF parasol RGCs, separately from electrical stimulation artifacts (see Methods). The resulting probability of a single, short-latency spike [21–23] as a function of current level for each cell was summarized with a sigmoidal activation curve (Fig. 1B, bottom). This electrical stimulus calibration thus revealed the set of electrodes and current levels that could selectively activate each target cell to fire a spike (Fig. 1B, right). Selective activation was defined as producing a spike in the target cell with probability >0.98 and in all other parasol cells with probability <0.02.

### Electrical stimulation replicates responses to natural images

The bidirectional electrical stimulus calibration was then used in the same experimental session to replicate the neural code at single-cell and single-spike resolution. First, 100 natural scenes jittered to simulate fixational eye movements were presented to the retina to identify naturalistic light responses (Fig. 2A, see Methods). The responses of the ON and OFF parasol RGCs to each image were identified from the MEA recordings. Then, for each image, a spatiotemporal sequence of electrical stimulation pulses chosen to reproduce the neural code in the two populations as faithfully as possible was determined: for each visually-evoked spike from each cell, a selective electrical stimulus for that cell was assigned at the corresponding time (Fig. 2B). Simultaneous stimulation of multiple cells using multiple electrodes was allowed when necessary. Stimuli were strategically chosen from the electrical stimulus calibration to mitigate sources of error. In particular, the selectively activating electrode nearest to the estimated soma center of each target cell reduced error resulting from minor physical displacement at the MEA-retina interface, and current amplitudes in the middle of the selective current range reduced error due to nearby multi-electrode stimulation (see Methods). The assigned spatiotemporal sequence of electrical stimulation was delivered to the retina and evoked spikes (as well as spontaneously-generated spikes) were identified in the recorded voltage traces (Fig. 2C, see Methods).

**Figure 2.**
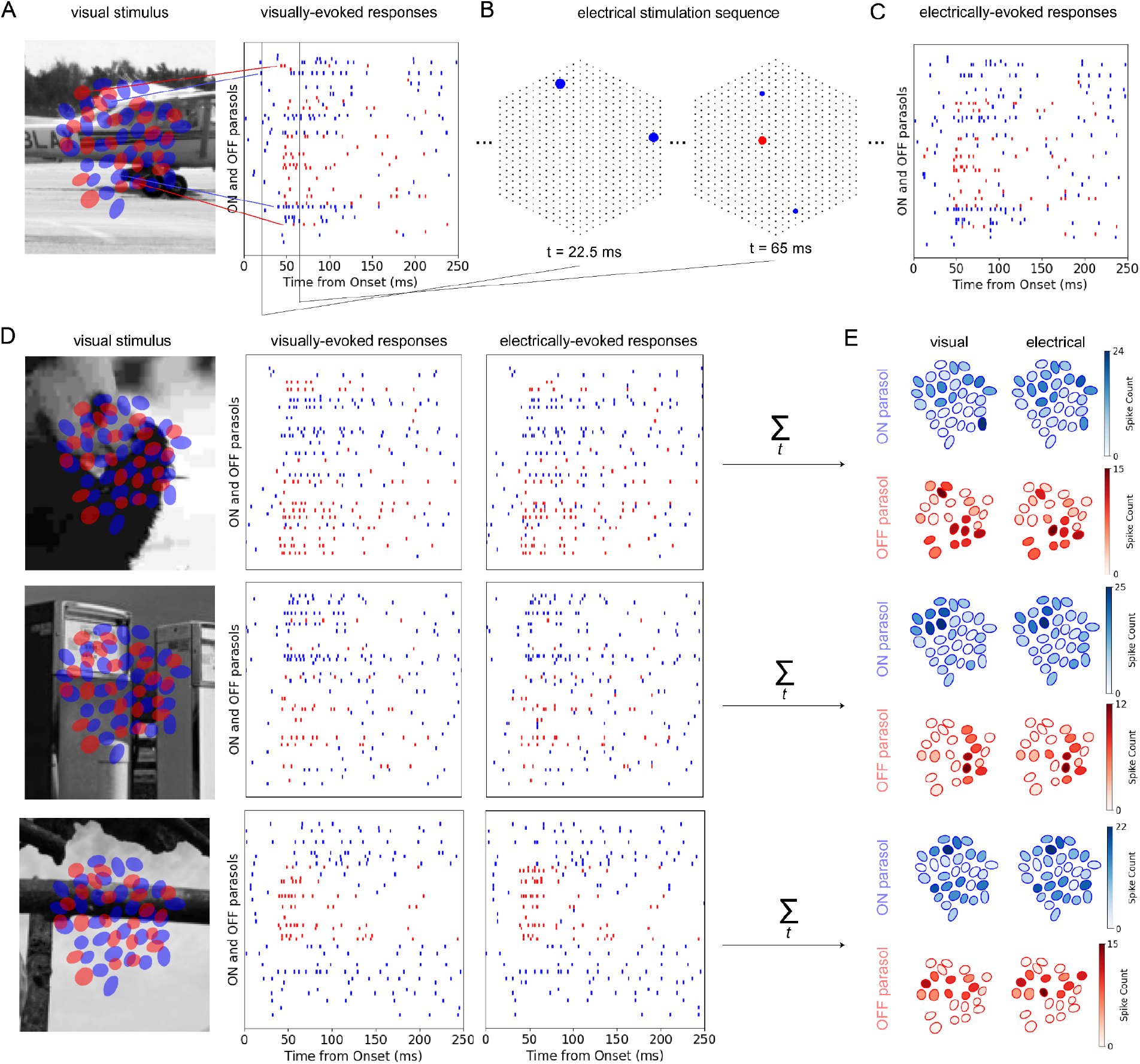
Reproducing naturalistic activity using electrical stimulation. (A) A naturalistic visual stimulus jittered to simulate fixational eye movements was presented to the retina, evoking responses in the ON (blue) and OFF (red) parasol RGC populations forming mosaics across the scene. RGCs were ordered in the spike raster plot according to RF location from top to bottom and colored according to their polarity. (B) An optimized electrical stimulation sequence was selected and delivered to faithfully reproduce the exact pattern of visually evoked responses from (A). The collection of black dots denote the electrode array. The collection of blue (red) points denote the stimulating electrodes at each time step, with the size of each point proportional to the current amplitude delivered and the color of each dot representing the polarity of the RGC being selectively targeted. (C) Electrically-evoked responses to the optimized electrical stimulation sequence, demonstrating precise encoding of the target neural activity in the intermixed ON and OFF parasol populations. (D) Three additional examples of reproducing naturalistic activity using electrical stimulation. In each case, the spatiotemporal electrical stimulation sequence was selected to faithfully reproduce the exact pattern of recorded responses to the visual stimulus. (E) Cell-type specific spatial encoding in the visually- and electrically-evoked responses. ON (OFF) parasol receptive fields are shaded blue (red) according to the total number of spikes recorded over the 250 ms. Note the complementary spatial encoding patterns between the oppositely-tuned neural populations present in the visually-evoked responses and maintained in the electrically-evoked responses.

Comparison of the visually-evoked responses to the electrically-evoked responses revealed reliable replication of the neural code for diverse natural images. Each image produced a distinct pattern of light-evoked activity in the two populations, and the electrically-evoked pattern of activity reproduced that light-evoked pattern with high fidelity (Fig. 2D). Notably, the highly differentiated spatial encoding patterns of the opposing ON and OFF cell types present in the visually-evoked responses were preserved in the electrically-evoked responses (Fig. 2E), despite the fact that the two populations were spatially intermixed, reproducing a fundamental aspect of the neural code.

### Electrical encoding is more precise than visual encoding

To quantify how reliably the natural activity of each cell was reproduced using electrical stimulation, responses to repeated trials of each naturalistic visual stimulus were compared with responses to repeated trials of the corresponding electrical stimulation sequence (Fig. 3A). The number and timing of action potentials fired by individual ON and OFF parasol cells were more variable across the visually-evoked response trials than across the electrically-evoked response trials. This trend was quantified using the Victor-Purpura spike train distance metric [24], which expresses differences between spike trains in terms of the elementary operations (spike shifting, deletion, and insertion) required to transform one spike train into another (see Methods). The average distance between the target visual response and visually-evoked responses in other trials was compared to the average distance between the target visual response and the electrically-evoked response trials, for each cell and for each visual image. The precision of the electrically-evoked responses was consistently higher than that of the natural visual code for the majority of the ON and OFF parasol cells (Fig. 3B).

**Figure 3.**
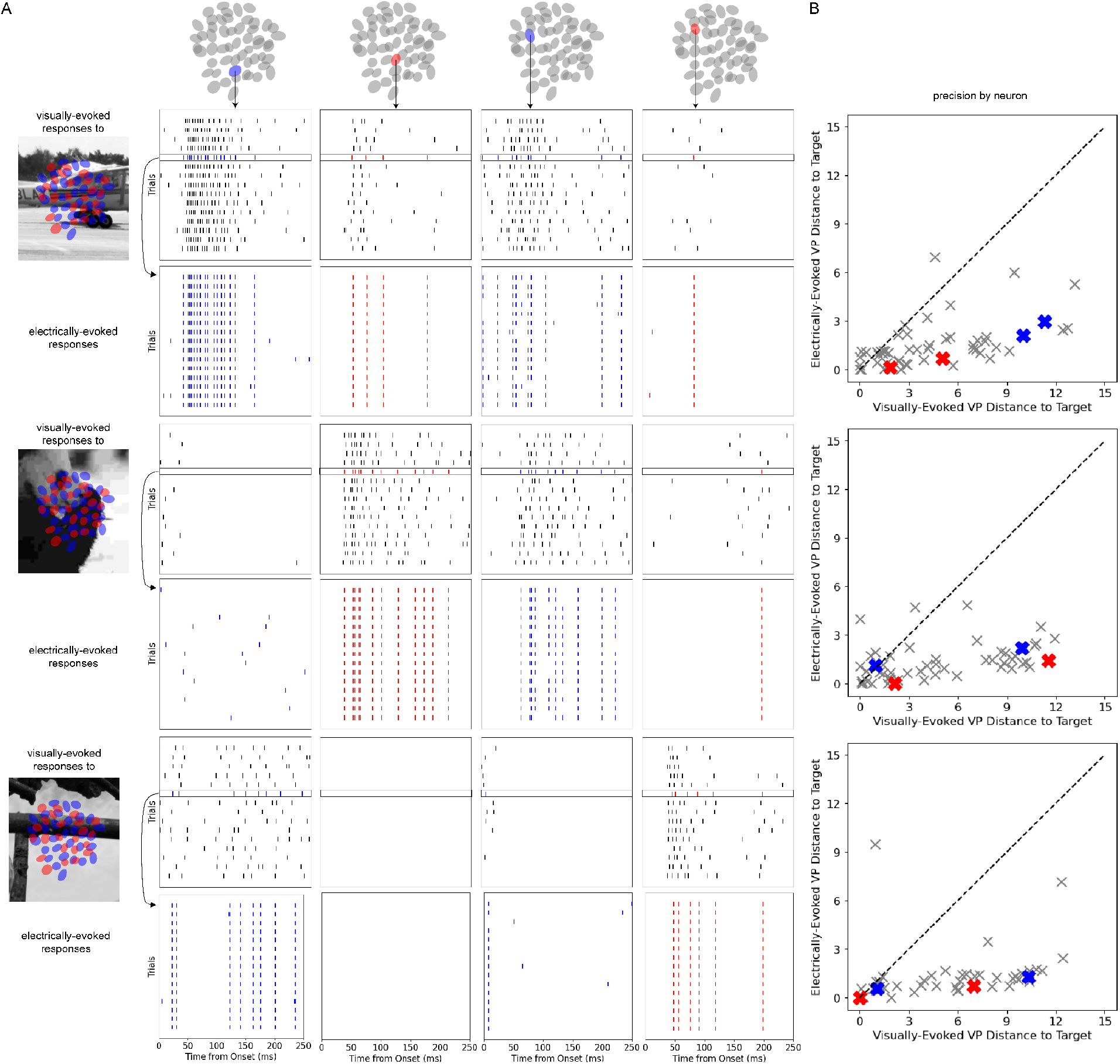
Spike train precision for individual cells in visually- and electrically-evoked responses. (A) 15 visually-evoked response trials to a naturalistic image and 15 electrically-evoked response trials to the optimized electrical stimulation sequence, for several representative ON and OFF parasol RGCs and for three naturalistic images. The visual response trial selected as target for each image is highlighted in color and encircled with a black box. (B) Scatter plots for each naturalistic image comparing the average Victor-Purpura distance between the target visual response and the other visually-evoked response trials versus the average distance between the target visual response and the electrically-evoked response trials for each individual parasol RGC. Featured RGCs from (A) are highlighted in the scatter plot.

To quantify spike train precision for the full ON and OFF parasol populations, distances between evoked population responses were determined by summing the Victor-Purpura distances obtained for each targeted cell (see Methods). This multi-neuron Victor Purpura distance was applied to all 15 visually-evoked and 15 electrically-evoked response trials associated with 5 natural images, generating a 150 by 150 matrix (Fig. 4A, top). Each 30 by 30 block along the main diagonal represents the distances (i.e. differences) between the 15 visual and 15 electrical stimulation trials for a given image, while the off-diagonal blocks represent the distances between responses associated with different images. As expected, visually- and electrically-evoked responses to a given image were more similar to each other than to those of the other images (30 by 30 blocks along the main diagonal were darker), verifying that distinct images were represented by distinct and repeatable neural code. Within each image block, the electrically-evoked responses were the most similar to the target visual response (dark horizontal and vertical lines) and to one another (dark 15 by 15 blocks along the main diagonal), verifying the enhanced ability of the electrical stimulation approach to reproduce the neural code in comparison with the natural visual code.

**Figure 4.**
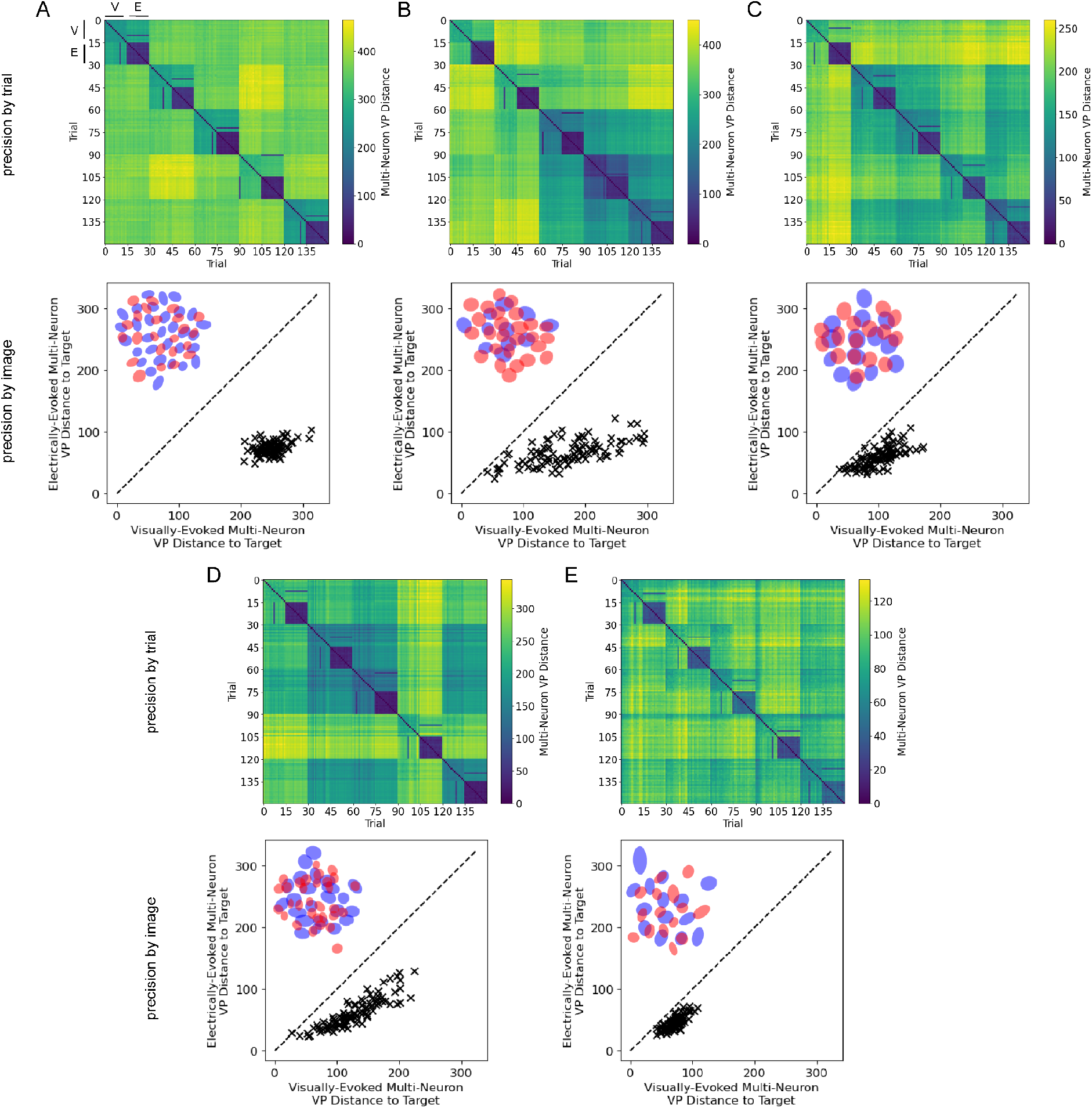
Spike train precision for complete populations in visually- and electrically-evoked responses in five retinas. (A) Spike train precision for the full ON and OFF parasol populations for the preparation from Figs. 1-3. Top: Multi-neuron Victor-Purpura distance for all 15 visually-evoked and 15 electrically-evoked response trials associated with 5 distinct natural images. “V” designates the visual stimulation trials and “E” designates the electrical stimulation trials for the first image; the distances between the 15 visual and 15 electrical stimulation trials for each individual image are the 30 by 30 blocks along the main diagonal. Bottom: Average multi-neuron Victor-Purpura distance from the target visual response to the other visually-evoked responses versus to the electrically-evoked responses for each of the 100 natural scenes. ON and OFF parasol RGC receptive field mosaics for the preparation are overlaid in the upper left quadrant. (B-F) Spike train precision for the full ON and OFF parasol populations for 4 additional preparations from 3 other macaque retinas, using the same natural images as for A.

The population spike train precision for each of the 100 natural images was summarized by comparing the similarity of the target visual response to the other visually-evoked responses, and to the electrically-evoked responses (Fig. 4A, bottom). Electrical stimulation evoked patterns of activity substantially more similar to the target neural code than repeated visual stimulation did for all 100 natural images. This trend was corroborated using an alternative means of assessing the spike trains: standard classification metrics of recall and precision applied to the full ON and OFF populations were much higher for electrically-evoked trials than for repeated trials of visual stimulation (see Methods, Extended Data Tab. 1).

The ability to reproduce the neural code was consistent across retinas. Across 5 total preparations from 4 macaque retinas, electrical stimulation more closely mimicked the target neural code than repeated visual stimulation did for 99% of natural images tested. On average across images and retinas, the activity evoked by electrical stimulation was 51 ± 17% more similar to the target than repeated visual stimulation. Similarly, the standard classification metrics of recall and precision were substantially higher with electrical stimulation for all recordings (Extended Data Tab. 1). Note that each preparation featured entirely distinct spatial organization of ON and OFF parasol RGCs relative to the electrode array (Fig. 4A-E). The consistency of spike train replication thus demonstrates the adaptivity of the approach: a bidirectional electrical calibration procedure rapidly reveals selective stimuli for the particular retina of interest, then an optimization procedure exploits that information within the same experimental session to reproduce population responses at their native resolution.

Unsurprisingly, electrical stimulation optimized only for parasol cells evoked nonspecific activity in other, non-targeted RGCs. To assess the impact of off-target stimulation, the evoked responses of 154 ON and OFF midget RGCs in the recording from Fig. 4A were analyzed. As expected, many of the electrically-evoked midget cell responses were substantially less similar to the target visual response than repeated responses to visual stimulation were to each other, and thus failed to replicate the neural code (Extended Data Fig. 1). This observation highlights the importance of closed-loop cell type targeting for restoring naturalistic patterns of activity, and the importance of targeting additional cell types in future work (see Discussion).

### Electrical stimulation replicates responses to moving objects

In addition to complex scene structure, natural vision involves moving objects, which evoke traveling waves of activity in parasol RGCs [19], including strongly localized activity at each time point which could be difficult to replicate with electrical stimulation. To test the ability to represent motion, a white moving bar on a uniform gray background was presented to the retina (Fig. 5A, see Methods) and RGC responses were recorded. Then, just as for the natural scenes, a corresponding spatiotemporal electrical stimulation sequence was determined and delivered to the retina, and evoked spikes (as well as spontaneously-generated spikes) were identified in the recorded voltage traces.

**Figure 5.**
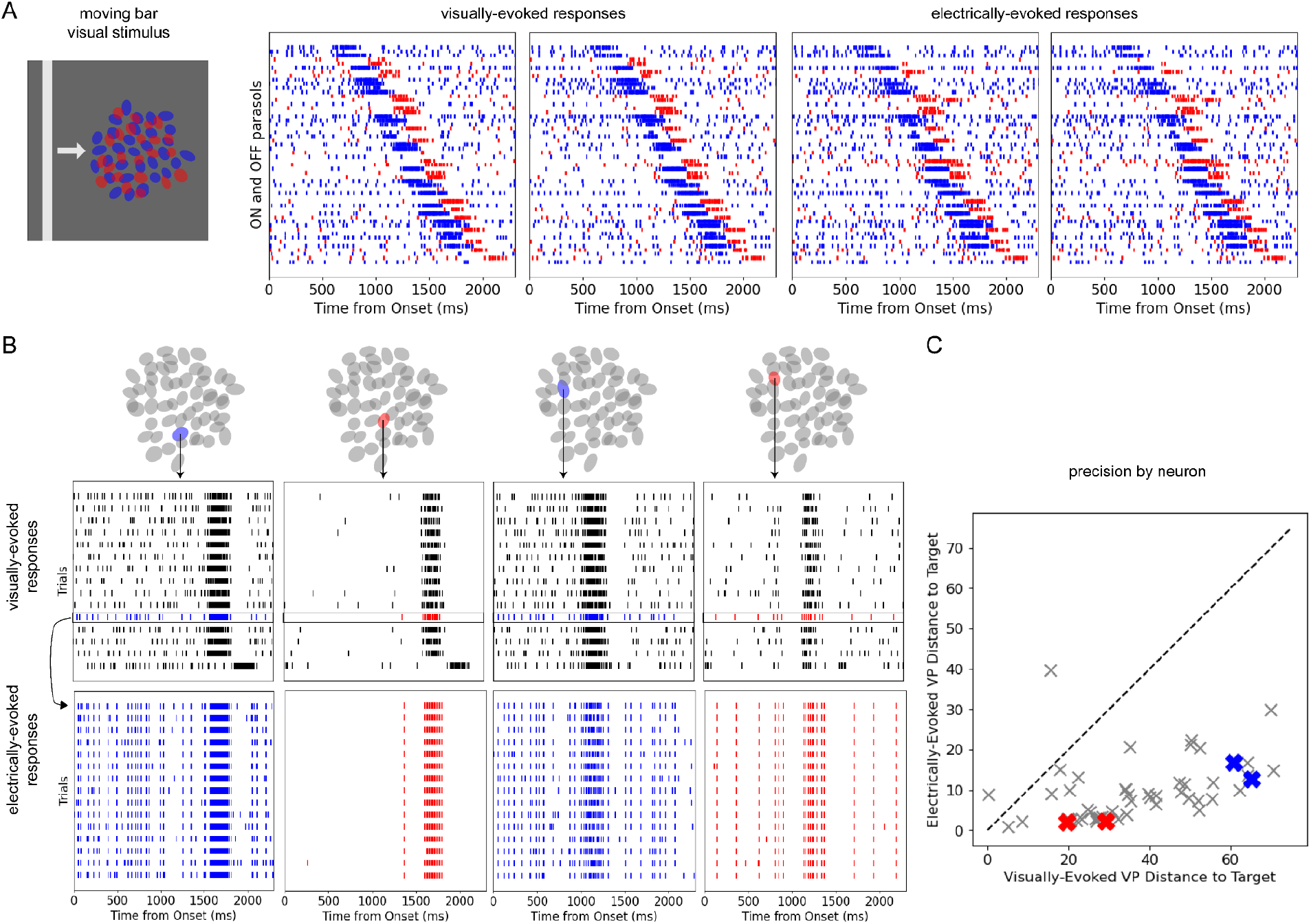
Reproducing moving bar activity using electrical stimulation. (A) Left: A moving bar visual stimulus was presented to the retina, evoking successive responses in the ON (blue) and OFF (red) parasol RGC populations forming mosaics across the scene. RGCs were ordered in the spike raster plot according to RF location from left to right and colored according to their polarity. Right: Several visually- and electrically-evoked RGC responses (2nd visually-evoked response is the target response for electrical stimulation). (B) All trials of visually- and electrically-evoked responses for several example ON and OFF parasol RGCs. (C) Scatter plot comparing the average Victor-Purpura distance between the target spike train and the other visually-evoked spike trains versus the electrically-evoked spike trains for each individual parasol RGC. Example RGCs from (B) are highlighted in the scatter plot.

Comparison of the electrically-evoked responses to the visually-evoked responses again revealed reliable replication of the neural code (Fig. 5A). The Victor-Purpura distance metric confirmed that electrical stimulation evoked patterns of activity more similar to the target visual responses than those evoked by repeated trials of visual stimulation, for the majority of cells recorded (Fig. 5B).

### Electrical stimulation enables arbitrary control of two neural populations

To investigate the potential of the bidirectional framework for applications beyond natural vision restoration, the same approach was used to produce structured patterns of neural activity in the parasol populations unrelated to the neural code of the retina. Artificial target patterns of activity were designed to mimic the appearance of words or images in the spike rasters, as a demonstration of an arbitrary manipulation of RGC activity. Sequences of electrical stimulation to replicate these artificial patterns were then delivered to the retina, as above. The resulting electrically-evoked spike patterns (combined with spontaneously-generated spikes) were encoded precisely enough to be legible or recognizable (Fig. 6), revealing the capacity of this framework for targeted scientific investigation or neural signal augmentation. The most obvious failure mode involved attempts to evoke many repeated spikes from the same neuron with less than ~10 ms between electrical stimuli (not shown) – unsurprisingly, the results in this extreme case were inconsistent, most likely due to spike refractoriness of the targeted cells.

**Figure 6.**
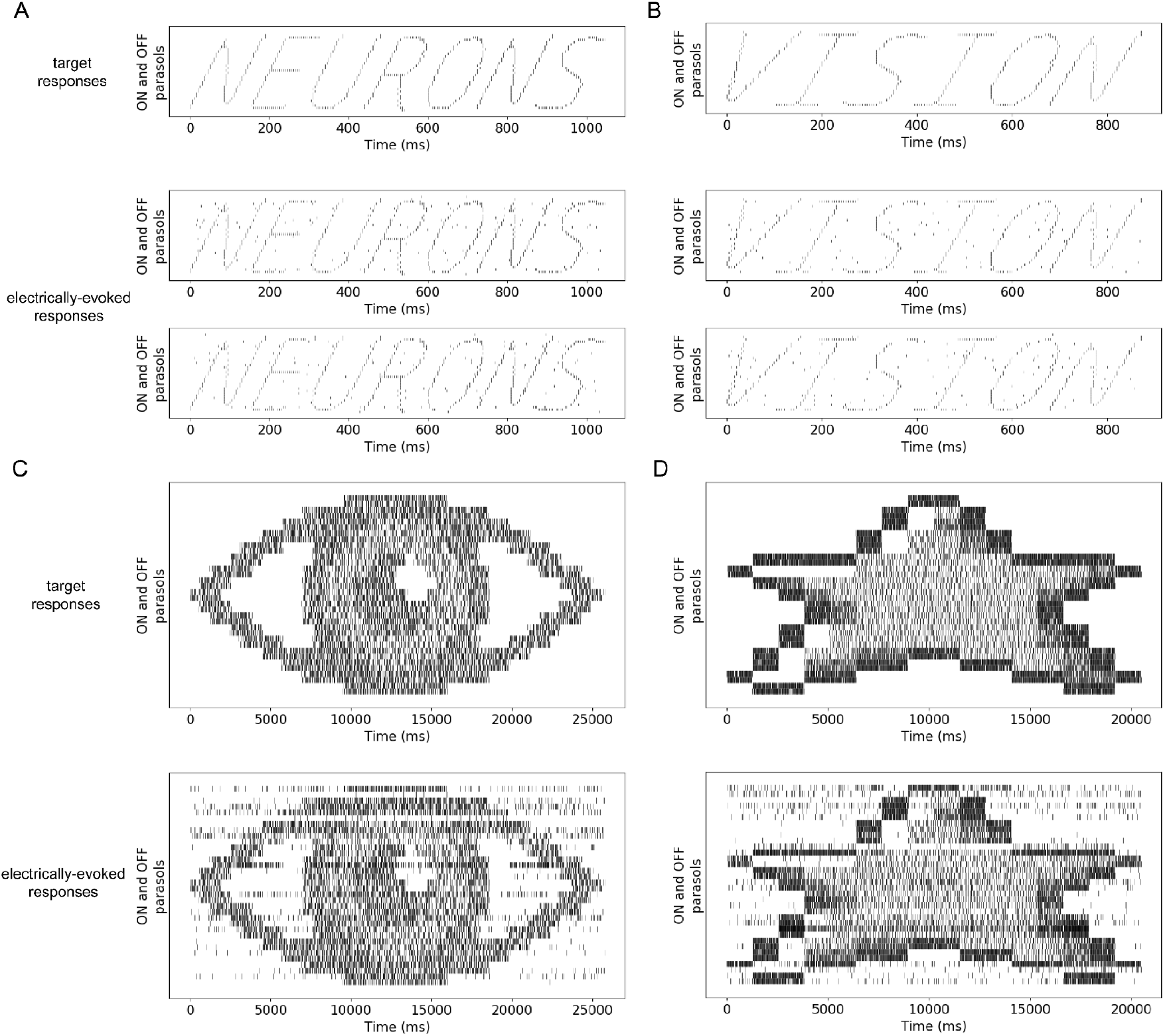
Arbitrary, structured control of neural responses. (A-B) Top: Artificial target patterns for ON and OFF parasol RGCs designed to invoke the appearance of words in the spike rasters. Bottom: Several trials of electrically-evoked responses corresponding to the above word targets. (C-D) Top: Artificial target patterns for ON and OFF parasol RGCs designed to invoke the appearance of images in the spike rasters. Activity for each parasol cell was divided into 320 ms bins and the spike rate per bin was modulated to invoke the appearance of different levels of shading in the spike rasters. Bottom: A trial of electrically-evoked responses corresponding to the above image targets.

## Discussion

Focal electrical stimulation replicated visually-evoked neural responses of intermixed populations of ON and OFF parasol cells at single-cell and single-spike resolution. The electrically-evoked activity demonstrated greater precision than natural visual responses, despite the dense and overlapping spatial organization of the two cell types. These results were consistent across retinas, indicating that the approach was robust to differences in the spatial arrangement of neurons in each retina. The approach was also successful in reproducing recorded traveling waves of activity in response to moving objects, and in evoking structured non-naturalistic activity patterns, raising the possibility of delivering novel visual signals to the brain. Together these results reveal the ability to control the activity of distinct neural populations simultaneously with greater fidelity than has been demonstrated previously, using a stimulation technology that is amenable to clinical translation.

### Relationship to prior work

Previous work highlights the potential impact of single-cell and cell-type specific electrical stimulation and the novel contribution of the stimulation approach used here. The ability to evoke individual spikes from primate RGCs with brief (~100 µs), weak (~1 µA) current pulses has been established in numerous studies [21–23]. Several studies demonstrated selective activation of individual neurons [21–23,25,26], and current steering using two or more stimulating electrodes has been shown to enhance spatial resolution [27–30]. Furthermore, an early proof-of-concept study demonstrated the ability to precisely reproduce the responses of six RGCs of one type to a moving bar visual stimulus, although this approach was not scalable and did not modulate multiple RGC types independently [31]. Another study presented an approach to optimize visual perception under partially selective activation of neural populations by choosing electrical stimuli which optimize reconstruction of the visual stimulus from the evoked responses, although this approach did not actually replicate the natural visual code [32]. Here, we extend this body of work toward practical and clinical application: faithfully replicating the responses of two large, complete and intermixed RGC populations to diverse naturalistic images and other stimuli, using a fully automated and scalable bidirectional approach.

The high spatial and temporal resolution achieved in this work is unique in the domain of vision restoration. Both epiretinal and cortical electrical stimulation approaches have typically treated electrodes as pixels, assigning each stimulating electrode to a fixed region of the visual field and representing this region by indiscriminately activating many neurons of multiple types near the electrode. The shapes and locations of these evoked percepts vary widely, across electrodes and across subjects, from round spots to elongated ellipses to lines and wedges. One likely contributor to this variability is the indiscriminate activation of nearby neurons of different types, including passing axons with cell bodies located far from the MEA [33]. Several different methods have been proposed and tested to leverage these percepts in useful ways via static or dynamic multielectrode stimulation strategies [34,35], but these methods are limited to simple patterns and do not extend to complex or complete visual scenes. To help address these limitations, the present approach used bidirectional calibration to identify and target individual cells, then reproduced their natural activity patterns to evoke an accurate representation of the neural code in those cell types.

A subretinal photovoltaic implant has made notable recent progress toward restoring form vision by interfacing with the retina at an earlier stage of visual processing — likely, the retinal bipolar neurons. Patients with this implant demonstrated the ability to read and write with letter acuity closely matching the 100 µm pixel pitch (20/420) [36]. The clinical performance of this implant is surprising given the indiscriminate nature of the electrical stimulation it provides: for example, simultaneous activation of ON and OFF RGCs would be expected to provide conflicting visual information to the brain. However, it is possible that indiscriminate extracellular stimulation of bipolar cells in the subretinal space produces greater ON RGC than OFF RGC activation due to inhibition in the retinal circuitry [37], or that the greater noise in OFF RGCs during degeneration [38] reduces the impact of evoked spikes on perception. Either of these possibilities could reduce the conflict between ON and OFF RGC signals. Furthermore, because the foveal area targeted in the photovoltaic approach is unique in being dominated by just two RGC types (ON and OFF midget), preferential activation of ON RGCs in the fovea could suffice for useful clinical performance. Regardless of the mechanism, these findings suggest that a reasonable degree of vision restoration is possible, in some conditions, without a technology that is designed to reproduce the neural code. Similarly, other neural interfaces have proven clinically useful for simple manipulations of neural activity without any attempt to reproduce the neural code [39–41]. In contrast, the approach in the present work is aimed at future high-fidelity functional restoration, in which reproducing the neural code precisely can hopefully improve the performance of interfaces to a variety of brain circuits.

Finally, a recent all-optical study combining two-photon optogenetic stimulation with two-photon calcium imaging demonstrated single-cell and sub-millisecond resolution reproduction of visually-evoked firing rates in dozens of neurons in the mouse brain [5]. While powerful for scientific investigation in some animal models, all-optical approaches present serious technical challenges for clinical translation: transgenic modification of neurons of each targeted type, expression and membrane trafficking of unnatural proteins, high intensity illumination, and optical hardware. Ultimately, these challenges reflect the technical overhead of transducing electrical signals into light (for optical recording), and light into electrical signals (for optical stimulation). In comparison, the present results achieve higher fidelity reproduction of the neural code in target populations using electrical stimulation and recording, a more direct form of reading and writing neural activity that is more conducive to translational application.

## Limitations

This study targeted only the ON and OFF parasol RGCs. As a result, while the approach precisely reproduced their spatiotemporal activity patterns, other RGCs were activated in non-naturalistic patterns (Extended Data Fig. 1). Previous data-driven simulations suggest that the resulting visual signal should remain perceptible despite the added high-spatial frequency noise from activation of the more numerous midget RGCs [42]. Therefore, the present approach is likely to be useful for vision restoration. However, given that parasol cells only account for approximately 20% of RGCs in the peripheral macaque retina [43], expanding the approach to consider the activation of additional RGC types will be an important next step.

Inadvertent activation of axon bundles arising from RGCs with somas located off the electrode array, which encode an unknown region of visual space, could also contaminate the artificial visual signal in a retinal implant [33]. In the present work, in some recordings, axon bundle activation due to electrical stimulation could be entirely avoided by using currents below axon bundle threshold; in others, at least one axon bundle was necessarily activated to achieve selective activation among complete parasol cell populations (Extended Data Tab. 1). Several factors likely contributed to this variation, including the retinal location targeted and the detailed local geometry of axon bundles. Strategies such as targeting the raphe region [42] or using penetrating electrodes to bypass the nerve fiber layer [44] may help to mitigate the impact of axon bundles.

Spontaneous activity could interfere with reliable encoding of information in RGCs. Indeed, this is one of the factors contributing to differences in electrical encoding precision among the different preparations (Fig. 4). Although spontaneous activity did not overwhelm the replication of visually-evoked spike trains in this study, higher spontaneous activity rates, which occur in at least some RGC types during degeneration [38], could significantly reduce the similarity of the electrical and visual encodings. Developing strategies to mitigate or compensate for high levels of spontaneous activity will be helpful for effective encoding.

The present results depended strongly on the ability to selectively stimulate individual ON and OFF parasol RGCs. This selectivity was achieved with electrode density ~10 times greater than the target cell density in the peripheral retina. However, in the central retina, where RGCs are much more densely packed for high-acuity vision, the achievable selectivity [42] and thus the ability to modulate two intermixed neural populations independently will be substantially reduced. Overcoming this challenge will likely require advances in electrode array design [44], improvements in current steering strategies [27–30], and/or more complex encoding algorithms [32].

Finally, restoring visual function after photoreceptor degeneration will require several additional steps beyond those shown in this work. Identification of neuronal types in the degenerated retina will require a different approach than the one used here, because light responses cannot be used. Recent work [45–47] has shown that electrical feature-based identification procedures can be used to reliably identify distinct cell types in the healthy primate retina, including ON and OFF parasol cells. However, more work is required to determine the effectiveness of these procedures in the degenerating retina. Additionally, target patterns of neural activity for the degenerated retina must be determined without recording natural responses to visual stimuli. A computational model of retinal processing that predicts RGC responses to visual stimuli [48–51] could be used to generate target activity patterns for electrical stimulation. The accuracy of the neural response model would then play a critical role in the success of the approach.

### Potential applications

The ability to precisely replicate the neural code with an electrical interface could support restoration of human vision, somatosensation, and other neurological functions. In addition to functional restoration, the ability to use electrical stimulation to evoke arbitrary, structured patterns of activity introduces new opportunities to investigate fundamental questions about how the neural code gives rise to computation, perception, and behavior. By enabling high-precision perturbation of neural activity across a broad range of neural systems, including in humans, electrical interfaces of this kind may provide a powerful foundation for understanding brain function and for the next generation of restorative neurotechnologies.

## Methods

### Multi-electrode array recordings

Retinas were obtained from terminally anesthetized macaque monkeys (*Macaca mulatta, Macaca fascicularis*) during the course of research performed by other laboratories, in accordance with Institutional Animal Care and Use Committee guidelines. Briefly, eyes were hemisected in room light immediately following enucleation and the anterior portion of the eye and vitreous humor were removed. The posterior portion of the eye was stored in darkness in oxygenated, bicarbonate-buffered Ames’ solution (Sigma-Aldrich) at 31-33° C.

Small (~2mm x 2mm) segments of retina with the attached retinal pigment epithelium (RPE) and choroid were separated from the sclera, after which the choroid was trimmed to improve tissue oxygenation. Segments in both temporal and nasal regions of the retina were obtained, with eccentricities ranging from 6-13 mm (27-41° temporal equivalent). The retina was mounted ganglion cell side down on a custom multielectrode array system with 512 electrodes (diameter 5-15 µm) arranged in an isosceles triangular lattice with center-to-center distance between adjacent electrodes in each row of 30 µm, covering a hexagonal area of 0.43 mm^2^ [52]. A transparent dialysis membrane was used to secure the retina against the array. During recording, the tissue was continuously superfused with oxygenated, bicarbonate-buffered Ames’ solution maintained at 33-35° C.

Raw voltage signals from the 512 channels were amplified, bandpass filtered (43-5000 Hz), multiplexed, digitized at 20,000 Hz, and stored for analysis, as described previously [53]. Spikes from individual RGCs in the recorded voltage traces were identified and sorted using standard spike sorting techniques [53].

### Visual stimulation

Visual stimuli were presented on a 120 Hz gamma-corrected cathode ray tube display (Dell UltraScan 800HS) and were optically reduced and projected through the mostly-transparent MEA onto the retina at low photopic light levels [54,55]. The visual stimulus area extended well beyond the recording area of the MEA.

For RGC type classification and receptive field mapping, a spatiotemporal white noise visual stimulus (flickering checkerboard) was presented for 15-30 minutes [56]. The stimulus refresh rate was either 30 or 60 Hz and the size of each stimulus pixel in the grid ranged from 36-102 µm at the photoreceptor layer. The intensity of each pixel was drawn randomly and independently from a binary distribution at each refresh. In some cases the three monitor primaries were modulated independently for spectral variation, and in other cases the three primaries were yoked for a black-and-white stimulus.

For evoking naturalistic responses with fixational eye movements, natural images from the ImageNet database [57] were used. Images were converted to grayscale values and displayed at 320 x 160 stimulus pixels, with each pixel measuring 9 or 17 µm at the photoreceptor layer. The average intensity across all images was 0.46 times the maximum, for all three display primaries. Images were displayed one after another in sequence for 500 ms each (60 frames at 120 Hz), with eye movements simulated as Brownian motion with a diffusion constant of 10 or 19 µm^2^/frame, similar to recorded fixational eye movements in both humans [58,59] and non-human primates (Z.M. Hafed and R.J. Krauzlis, personal communication, June 2008). The entire sequence of 100 images was repeated 15 times. To evoke traveling waves of activity, a moving bar stimulus was presented drifting across a uniform field (width 0.25 or 0.5°, speed 2.7 or 5.1 deg/s, background intensity 0.1-0.5, bar intensity 0.88-0.95), and was repeated 15 times.

### Cell type classification

The spike-triggered average (STA) from the white noise stimulus [56] was computed for each cell to reveal the spatial, temporal, and chromatic properties of the light response. Distinct RGC types were classified by identifying distinct clusters in the light response and spiking properties, including features of the time course and spike train autocorrelation function extracted via principal component analysis, and the spatial extent of the receptive field [18–20,60,61]. All recordings included cells of the four numerically dominant types in primates: ON parasol, OFF parasol, ON midget, and OFF midget. In most preparations, recorded populations of ON and OFF parasol cells formed nearly complete mosaics over the region of retina recorded, whereas recorded populations of ON and OFF midget cells were less complete. An elliptical approximation of the spatial receptive field extent for each cell was obtained by fitting a 2D Gaussian to the STA frame with the maximum signal amplitude. The 2σ contour of these fits is used to illustrate the mosaic organization.

### Electrical stimulation

Electrical stimulation was delivered through one or more electrodes while recording RGC responses from all electrodes simultaneously. For the entire duration of all electrical stimulation, a uniform gray screen was presented on the display and projected onto the retina. Two types of electrical stimulation patterns were used: single-electrode stimulation for electrical stimulus calibration and variable-electrode stimulation for reproducing recorded activity patterns.

To perform the electrical stimulus calibration, a sequence of single-electrode stimuli were delivered to the retina. Each stimulus consisted of a charge-balanced, triphasic current pulse passed through an individual electrode on the MEA. The triphasic pulse was composed of anodal/cathodal/anodal phases with relative current amplitudes 2:-3:1 and pulse duration of 50 µs/phase (150 µs total), chosen to minimize the electrical artifact [23,25,52]. The current amplitude range tested (second phase) was 0.1-4 µA (42 amplitudes, linear scale), with each amplitude repeated 15 times. One stimulus was delivered every 10 ms in a pseudo-random order, restricted so that each successive stimulating electrode was far from the previous and subsequent electrodes, to avoid stimulating the same cells within their refractory periods.

To reproduce neural population activity, spatiotemporal sequences consisting of variable numbers of electrical stimuli were delivered to the retina, depending on the target activity pattern. For each visual stimulus, the response trial with the median number of total spikes in the ON and OFF parasol populations was designated as the target activity pattern. The spikes from the target were rounded to the nearest 2.5 ms, to guarantee enough time between electrical stimuli to facilitate spike sorting in the presence of electrical stimulation artifacts. Thus, spikes in the rounded template were shifted in time by a maximum of 1.25 ms relative to their true timing. For each spike from each cell in the rounded template, a selective electrical stimulus for that cell was assigned at the corresponding time. Therefore, each spatiotemporal electrical stimulation sequence consisted of 0 or more simultaneous stimuli every 2.5 ms, depending on the presence of target spikes during each of those periods. Across all natural scenes for the featured retina, there were on average 2.3 stimulating electrodes per pattern, with average electrode separation 194 µm. For the moving bar, there were on average 2.1 stimulating electrodes per pattern, with average separation 193 µm.

The selective electrical stimuli used during this assignment procedure were identified by the electrical stimulus calibration, which revealed the set of electrodes and current levels that could selectively activate each target cell. In principle, any of these selective stimuli could be used to activate the target cell without activating any non-target ON or OFF parasol RGCs. In practice, however, stimuli were strategically chosen from the electrical stimulus calibration to mitigate sources of error. First, to compensate for any physical displacement at the MEA-retina interface during the experiment, the selective electrode nearest to the estimated soma center of each target RGC was used. The soma center of each RGC was estimated as the spatial average of the locations of all electrodes recording somatic waveforms from the cell, weighted according to the maximum amplitude deflection at each electrode. Second, note that nearby simultaneous stimuli (<350 µm) could potentially induce deviations from single-electrode calibrated responses [32]. To increase tolerance to these deviations, the current amplitude for each stimulating electrode was chosen to bisect the selective current range, thus allowing limited movement of the target and non-target sigmoidal activation curves without compromising selectivity.

### Identifying responses to electrical stimulation

Electrical stimulation produced large, rapidly decaying artifacts that could obscure the directly-evoked sub-millisecond latency RGC spikes. To identify stimulus-evoked spikes efficiently, we used two complementary spike-sorting approaches introduced in accompanying work [62]. One approach leveraged repeated presentations of an identical stimulus to separate spiking and non-spiking cases by clustering post-stimulus recordings (“GraphSort”), and a second approach enabled single-trial detection by matching known spike templates to artifact-free portions of the recorded waveform (“PartialSort”). GraphSort was applied to the electrical stimulus calibration and to activity pattern reproduction; PartialSort was independently applied to activity pattern reproduction where possible.

For the single-electrode stimulation used to calibrate responses at each of the 512 electrodes across a range of current amplitudes, electrically-evoked spikes from RGCs were identified using GraphSort. For each electrode and current amplitude, post-stimulus voltage traces from 15 repeated trials were collected and restricted to electrodes expected to contain an informative signal for each cell based on its electrical image (EI) — typically electrodes with somatic waveforms and high signal-to-noise ratio. Traces were then clustered across repeats (using affinity propagation followed by distance-based merging), and pairwise cluster difference waveforms were computed after alignment to isolate the incremental contribution of spikes. These difference waveforms were then compared against EI templates in all candidate neurons and for a range of latencies to identify the previously-recorded cell whose spike waveform best explained each cluster transition. This procedure yielded an estimated spike probability for each electrode-cell pair at each current amplitude. Response probability as a function of current amplitude was summarized by fitting a sigmoidal activation curve over current amplitudes for each electrode-cell pair of the form 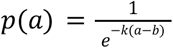, where *p*(*a*) is the probability of a spike, *a* is the current amplitude, and *k* and *b* are free slope and threshold parameters, using maximum likelihood estimation. The fitted parameters for every electrode-cell pair were subsequently used to identify selective stimuli.

For the spatiotemporal electrical stimulation sequences used to reproduce target activity patterns, electrically-evoked spikes from RGCs were identified using GraphSort and, independently, using PartialSort. Each stimulation sequence was delivered 15 times to facilitate the use of GraphSort.

First, GraphSort was applied to the post-stimulus voltage traces collected from each trial for each electrical stimulus. For cases in which all trials formed a single cluster, preventing the assignment of candidate neurons to difference waveforms, responses to single-electrode stimulation obtained both before and after the stimulation sequence were used to disambiguate spike probability for each cell between 0 or 1. Spontaneously-generated spikes from RGCs in the remainder of the recorded voltage traces were identified and sorted using standard spike sorting techniques [53]. This GraphSort approach was most reliable for identifying responses to effectively independent stimuli with sufficient separation in space and time. However, the GraphSort approach could be susceptible to errors in cases involving interactions between stimulating electrodes which evoked deterministic activity. In the first case, the GraphSort approach could be subject to false negatives, since it could miss deterministic spiking of off-target cells which did not spike in response to any of the individual stimuli. In the second case, the GraphSort approach could be subject to false positives, since it could miss deterministic suppression of a target cell which did spike in response to one of the individual stimuli. The latter case is much less likely because additional current injection beyond the single-electrode stimulation delivered during the calibration is more likely to cause another cell to fire than to suppress firing [63].

Second, PartialSort was also applied to the voltage traces collected from each of the 15 repeated trials for each stimulation sequence. Electrically-evoked spikes were detected on individual trials by template matching to the artifact-free portion of the post-stimulus recording. In this approach, samples in an early post-stimulus window dominated by residual artifact were excluded, and the remaining voltage was explained as a superposition of EI templates at candidate latencies using a greedy matching procedure; detections were accepted only when a sufficient fraction of the template overlapped artifact-free samples and the match exceeded a noise-relative threshold. Spontaneously-generated spikes were detected in the remainder of the recorded voltage traces using the same template matching approach. The PartialSort approach was useful for identifying responses to complex spatiotemporal stimuli for which constructing stable “spike-present” versus “spike-absent” comparisons was impossible due to deterministic spiking. However, by directly considering the voltage traces, the PartialSort approach could be subject to false positives, due to voltage deflections from extraneous sources such as axon bundle activation and due to necessarily lower template matching thresholds. For this reason, PartialSort was only applied to experiments delivering electrical stimulation exclusively below the axon bundle threshold (Extended Data Tab. 1). Under these conditions, results from both the GraphSort and PartialSort approaches could be compared directly to bound performance.

### Quantification of spike train similarity

To quantify the precision of the artificially evoked neural code for individual cells, the Victor-Purpura spike train distance metric [24] was computed between the target visual trial and repeated visual trials and between the target visual trial and electrical trials. The metric expresses distance between spike trains in terms of the cost of elementary operations (spike shifting, deletion, and insertion) required to transform one spike train into another: inserting or deleting a spike costs 1.0 and shifting a spike by an amount ∆t costs q * ∆t. Thus, the cost parameter q adjusts the timescale of comparison: small q emphasizes rate coding, while large q emphasizes temporal coding. At q=0, the metric equals the absolute difference in spike counts, and increasing q increases the sensitivity to precise spike timing. The results presented in this work used q = 1/(5 ms), reflecting results from previous studies investigating the temporal resolution of visual signals [64]. However, the main results were consistent across a larger parameter range (Extended Data Fig. 2).

To quantify the precision of the artificially evoked neural code for the full ON and OFF parasol populations, the Victor-Purpura distances obtained for each cell were summed. This corresponds to the standard multi-neuron Victor-Purpura spike train distance metric for a labeled-line code [65], as appropriate for the retina. Briefly, the multi-neuron Victor-Purpura metric adds a cost k for changing the neuron label of a spike. However, in the case of the retina, since each RGC sends unique information about a specific location in the visual field to distinct target(s) in the brain, we treat the responses as a labeled-line code and therefore the neuron label of a spike should never be changed. Therefore, we set k=2 to prevent changing the neuron label, and the multi-neuron Victor-Purpura distance between two population responses reduces to the sum of the Victor-Purpura distances between the pair of responses for each neuron.

Summary multi-neuron Victor-Purpura distances for each experiment were pooled across trials and images and reported as mean ± standard deviation (Extended Data Tab. 1). A second comparison method was used to independently corroborate the reproduction performance of the artificially evoked neural code. Standard classification metrics of precision and recall were computed using the target visual trial as ground truth. Specifically, recall was defined as the fraction of spikes from an electrical trial matching the target visual trial (i.e., a spike from the same cell within ±2.5 ms) out of the total number of spikes in the target visual trial. Precision was defined as the fraction of spikes from an electrical trial matching the target visual trial out of the total number of spikes in the electrical trial. Summary precision and recall metrics for each experiment were pooled across trials and images and reported as mean ± standard deviation (Extended Data Tab. 1).

### Generation of artificial target patterns

Artificial target patterns were designed to invoke the appearance of words or images in spike rasters as a demonstration of arbitrary manipulations of neural activity. All parasol RGCs whose assigned selective electrode belonged to the outer two rings of electrodes on the MEA were excluded from the spike rasters to avoid potential calibration errors at the MEA boundary, the most likely location for the MEA-retina interface to undergo physical displacement.

For the word patterns, spike rasters were derived from text graphics. Sequential spikes from the same cell were separated by 5 ms or more to avoid repetitive refractory period violations in the artificial target pattern. On average across 8 word patterns for the featured retina, there were 2.5 stimulating electrodes per pattern, with average electrode separation 239 µm.

For the image patterns, spike rasters were derived from images. The 2D array of grayscale intensities representing the image was used to generate differential spike rates across cells and time bins. Briefly, each row of the 2D array was assigned to a cell, and each intensity value in the row specified the spike rate for that cell in the corresponding time bin. A number of spikes proportional to the intensity value (between 0-8 spikes) was allocated within each time bin (320 ms), modulating the spike rate between 0-25 Hz. Spikes were evenly assigned throughout each time bin, then jittered in time by random samples from a zero-mean Gaussian distribution with σ=160 ms/x where x was the number of spikes allocated in that time bin. On average across 2 image patterns, there were 2.5 stimulating electrodes per pattern, with average electrode separation 201 µm.

---

**Extended Data Table 1.** Spike train reproduction metrics across retinas. Victor-Purpura distance, recall, and precision are computed for the full ON and OFF parasol populations across all electrical trials and images and are reported as mean ± standard deviation. PartialSort (blue) was run independently of GraphSort (black) as a means to bound results where possible. For comparison, the same metrics were also computed across all repeated visual trials and images (gray). The experiment featured in the figures is designated with an asterisk. The set of experiments featured in Figure 4 are from the standard condition. Bold boxes aggregate data across retinas.

|  | retina | multi-neuron VP distance | recall | precision |
| --- | --- | --- | --- | --- |
| standard | 1* | 74 ± 10<br>248 ± 19 | 0.91 ± 0.02<br>0.27 ± 0.07 | 0.85 ± 0.04<br>0.27 ± 0.07 |
|  | 2 | 67 ± 20<br>171 ± 60 | 0.92 ± 0.04<br>0.36 ± 0.09 | 0.79 ± 0.09<br>0.36 ± 0.09 |
|  | 3 | 60 ± 16<br>102 ± 28 | 0.82 ± 0.07<br>0.26 ± 0.11 | 0.64 ± 0.10<br>0.27 ± 0.11 |
|  | 4 | 62 ± 24<br>127 ± 41 | 0.78 ± 0.09<br>0.16 ± 0.12 | 0.66 ± 0.09<br>0.17 ± 0.13 |
|  | 5 | 46 ± 11<br>71 ± 13 | 0.64 ± 0.09<br>0.19 ± 0.08 | 0.57 ± 0.09<br>0.20 ± 0.08 |
|  | **all** | **62 ± 19**<br>**144 ± 72** | **0.81 ± 0.12**<br>**0.25 ± 0.12** | **0.70 ± 0.13**<br>**0.25 ± 0.12** |
| electrical stimulation below axon bundle threshold | 1 | 69 ± 9<br>97 ± 12<br>211 ± 20 | 0.83 ± 0.04<br>0.87 ± 0.03<br>0.27 ± 0.07 | 0.86 ± 0.05<br>0.72 ± 0.06<br>0.27 ± 0.07 |
|  | 2 | 67 ± 18<br>225 ± 50<br>153 ± 53 | 0.86 ± 0.08<br>0.91 ± 0.05<br>0.36 ± 0.09 | 0.77 ± 0.11<br>0.42 ± 0.09<br>0.36 ± 0.09 |
|  | 3 | 59 ± 15<br>60 ± 11<br>90 ± 25 | 0.62 ± 0.11<br>0.85 ± 0.08<br>0.27 ± 0.11 | 0.62 ± 0.14<br>0.60 ± 0.11<br>0.27 ± 0.11 |
|  | 4 | 32 ± 13<br>82 ± 29<br>78 ± 28 | 0.75 ± 0.08<br>0.84 ± 0.08<br>0.17 ± 0.14 | 0.74 ± 0.13<br>0.40 ± 0.08<br>0.18 ± 0.15 |
|  | 5 | 47 ± 15<br>92 ± 19<br>63 ± 13 | 0.56 ± 0.12<br>0.76 ± 0.08<br>0.18 ± 0.08 | 0.52 ± 0.12<br>0.32 ± 0.06<br>0.19 ± 0.09 |
|  | **all** | **55 ± 20**<br>**111 ± 65**<br>**119 ± 63** | **0.72 ± 0.15**<br>**0.85 ± 0.08**<br>**0.25 ± 0.12** | **0.70 ± 0.16**<br>**0.49 ± 0.17**<br>**0.25 ± 0.12** |

**Extended Data Figure 1.**
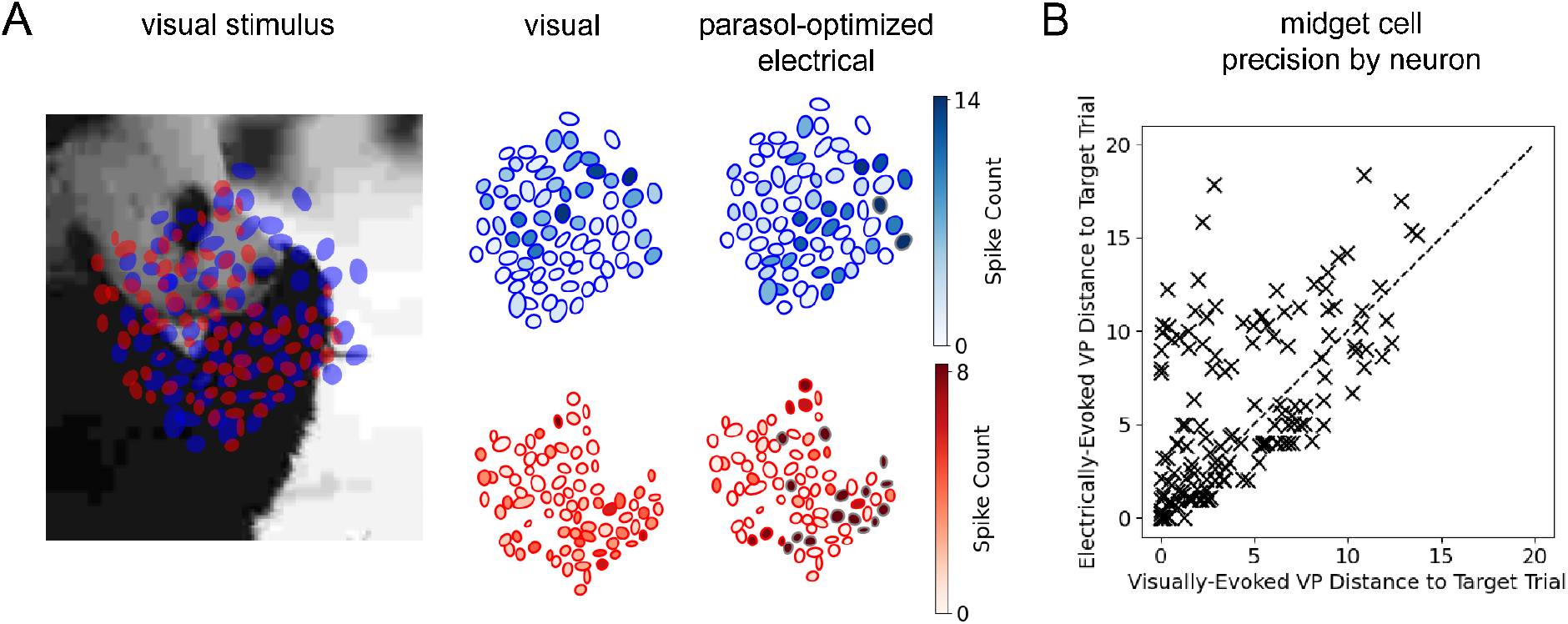
Impact of parasol-optimized electrical stimulation on midget retinal ganglion cells. (A) Cell-type specific spatial encoding in ON and OFF midget populations resulting from visual stimulation with a jittered natural scene and the corresponding parasol-optimized electrical stimulation. ON (OFF) midget receptive fields are shaded blue (red) according to the total number of spikes recorded over 250 ms, averaged across trials. Spike counts exceeding the visual response range are designated by gray receptive field outlines. (B) Scatter plot comparing the average Victor-Purpura distance between the target visual response trial and the other visually-evoked response trials versus the average distance between the target visual response trial and the parasol-optimized electrically-evoked response trials for each individual midget RGC.

**Extended Data Figure 2.**
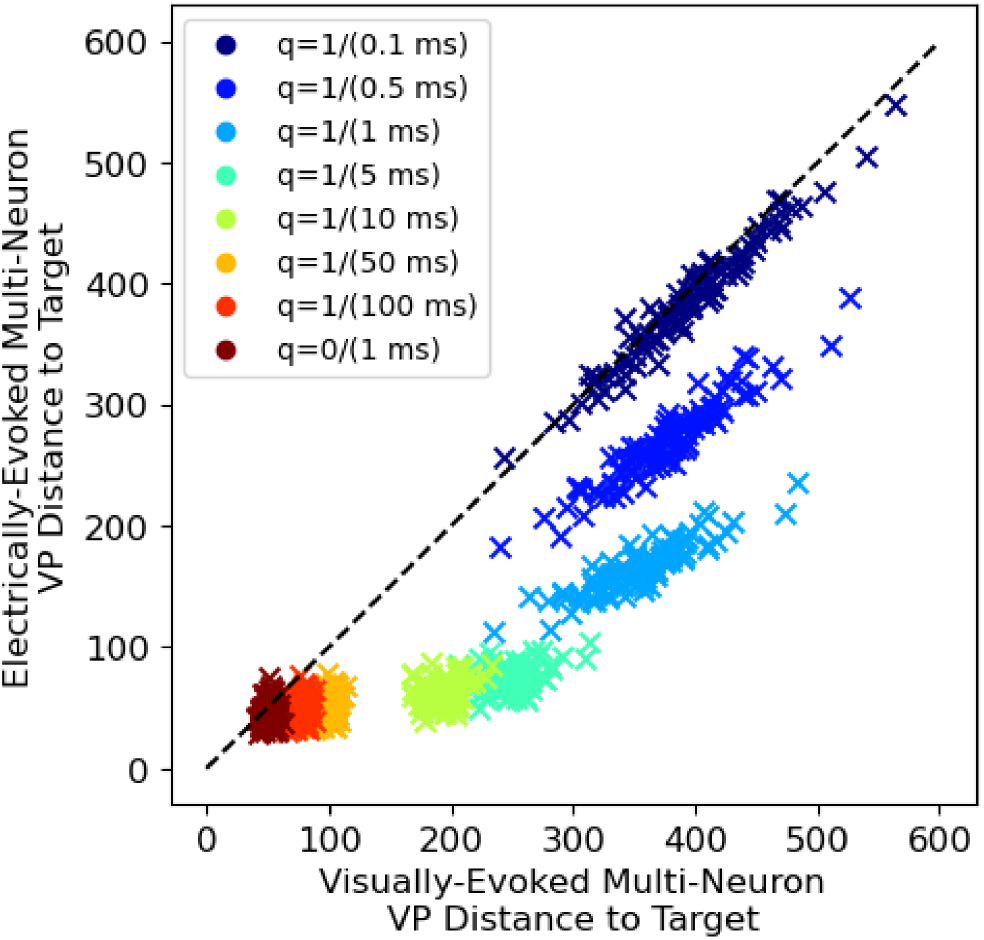
Spike train precision for Victor-Purpura cost parameter sweep. Average multi-neuron Victor-Purpura distance from the target visual response to the other visually-evoked responses versus to the electrically-evoked responses for each of the 100 natural scenes, for different values of Victor-Purpura cost parameter q. q=1/(5 ms) was used for the results presented in this work.

## Acknowledgements

This research was supported by National Science Foundation Graduate Research Fellowship Grant No. 2146755 and NSF Grant No. 1828993 (A.J.P.), Stanford Bio-X SIGF Fellowship and Stanford NIH T32 Biotechnology Training Grant (A.L.), NEI grants R01-EY021271 and R01-EY032900, NIH Blueprint MedTech through NEI and NIBIB grant U54-EB033650, and Wu Tsai Neurosciences Institute (E.J.C.). We thank T. Moore for access to primate retinas; N. Shah, S. Madugula, E. Wu, S. Cooler, R. Vilkhu, A. Gogliettino, G. Field, and S. Mitra for useful discussions; and K. Affolder, S. Kachiguine, D. Lopes, and S. Cital for technical assistance.

## Author Contributions

A.J.P., A.L., and E.J.C. designed research; A.J.P., A.K., M.H., P.V., B.H., A.S., M.A.S., C.B., Y.L., and E.J.C. performed multi-electrode array experiments; A.J.P., A.L., P.V., B.H., J.B., and H.N. contributed analytic tools, A.J.P., A.K., and E.J.C. analyzed the data; P.H., A.S. and A.M.L. provided and supported multi-electrode array technology; A.J.P. and E.J.C. wrote the manuscript.

